# A role for YAP-mediated regulation of invadopodia in HNSCC cells

**DOI:** 10.1101/2024.08.02.606435

**Authors:** Rachel J. Jerrell, Dayton D. Marchlewski, Aron Parekh

## Abstract

The objective of this study was to determine whether nuclear translocation of the transcriptional coactivator Yes-associated protein (YAP) was sensitive to extracellular matrix (ECM) rigidity and promoted Rho-associated kinase 2 (ROCK2) expression to affect invadopodia maturation and ECM degradation. ECM rigidity mimicking head and neck squamous cell carcinoma (HNSCC) tumor mechanical properties was simulated in vitro using a well-established model based on fibronectin-conjugated polyacrylamide gels (PAAs). The ratio of nuclear to cytoplasmic YAP and overall cellular ROCK2 levels were evaluated in HNSCC cells using quantitative immunofluorescence. YAP-mediated ROCK2 expression in HNSCC cells was determined using nested PCR and Western blot in response to the YAP inhibitor verteporfin. Invadopodia and ECM degradation were evaluated in HNSCC cells with siRNA-mediated inhibition of YAP using quantitative immunofluorescence in invadopodia assays. Both YAP nuclear translocation and ROCK2 cellular levels increased with ECM rigidity. Inhibition of YAP activity with verteporfin decreased ROCK2 gene and protein expression. Knockdown of YAP with siRNA inhibited the formation of mature invadopodia and ECM degradation but not total invadopodia (i.e., mature and immature or not degrading). Our study suggests that tumor-associated ECM rigidity can promote mechanically-induced transcriptional regulation to control proteolytic activity by affecting invadopodia maturation.

## 1. INTRODUCTION

The spread of migrating cancer cells from the primary tumor site requires their invasion through the extracellular matrix (ECM) of neighboring tissues.^1^ Cell migration is regulated by actomyosin contractility via phosphorylation of the myosin light chain (pMLC) of non-muscle myosin-II (NM II) by several kinases, including Rho-associated kinase (ROCK), in a process that is promoted by tumor-associated ECM rigidity known to induce malignant cellular behavior.^2-4^ Rigidity-dependent actomyosin contractility can activate several transcriptional programs including those regulated by the Yes-associated protein (YAP) which translocates into the nucleus to coordinate the expression of target genes with the TEA domain (TEAD) family of transcription factors.^5-7^ YAP is emerging as a key intracellular mechanotransducer, and YAP levels have recently been found to correlate with HNSCC tumorigenesis, progression, and poor prognoses, but the mechanistic role of YAP in cellular invasion is unknown.^8-13^

To breach dense tissues, migrating cancer cells typically utilize actin-rich adhesive protrusions called invadopodia that localize proteinases to create paths by focally degrading the ECM to facilitate tissue invasion.^14-16^ Invadopodia dynamics are driven by actin assembly at the cell membrane that promotes the elongation of these subcellular structures necessary for stable and mature degrading protrusions.^17-19^ We previously established that cellular traction forces generated by NM II-dependent actomyosin contractility in response to ECM rigidity correlate with ECM degradation by invadopodia in HNSCC cells.^20^ However, we later discovered that force generation and invadopodia maturation in HNSCC cells were differentially regulated by the ROCK isoforms, ROCK1 and ROCK2.^21^ Specifically, we found that ROCK1 mediates the traction forces necessary for migration by regulating actomyosin contractility via pMLC while ROCK2 promotes the maturation of invadopodia into actively degrading structures downstream of these forces by affecting local actin polymerization at the cell membrane via LIM kinase (LIMK).^21^

Recently, YAP levels have been found to correlate with ROCK2 expression in mesothelioma tumors, and YAP was also identified as a rigidity-dependent transcriptional regulator of ROCK2 via actomyosin contractility in a human breast cancer line.^22,23^ Since actomyosin contractility can facilitate YAP nuclear translocation, we hypothesized in this study that ECM rigidity can regulate YAP activity to drive ROCK2 expression thus mediating invadopodia maturation and ECM degradation by HNSCC cells.

## 2. MATERIALS AND METHODS

### 2.1 Cell culture and reagents

SCC-61 cells, an invasive HNSCC cell line originally acquired from the Yarbrough laboratory at our institution, have served as an established cell line model for invadopodia in our laboratory and were cultured as previously described (available from MilliporeSigma, Cat# SCC280, RRID: CVCL_7118).^20,21,24,25^ Chemical inhibition of YAP was performed using verteporfin (Sigma) which inhibits YAP/TEAD interactions.^26,27^ Kockdown (KD) of YAP was performed using a single ON-TARGETplus siRNA (sequences available from Dharmacon).

### 2.2 Nested PCR

RNA was extracted from cells using Qiagen RNeasy kit (Qiagen) per the manufacturer’s instructions. cDNA was generated using the Bio-Rad iScript cDNA kit (Bio-Rad) per the manufacturer’s instructions. ROCK2 human primer set A was used to amplify the PCR product which was amplified again using set B (Santa Cruz) per the manufacturer’s instructions. The amplified product was then run on a 1% agarose gel and visualized with SYBR Safe DNA Gel Stain (ThermoFisher) per the manufacturer’s instructions.

### 2.3 Western blotting

Western blotting for ROCK2 and YAP were performed as previously described using an anti-ROCK2 mouse monoclonal antibody (Abcam, Cat# ab-56661, RRID: AB_945286) and anti-YAP mouse monoclonal antibody (Santa Cruz, Cat# sc-101199, RRID: AB_1131430).^21^

### 2.4 Invadopodia assays with polyacrylamide gels

Invadopodia assays were performed using polyacrylamide gels (PAAs) that provide normal and malignant tissue mechanical properties as previously described.^2,20,28-30^ Briefly, fibronectin-conjugated soft and hard PAAs with elastic moduli of 1,023 and 7,307 Pa, respectively, were cast on glass bottom dishes (Mattek). PAAs were overlaid with cross-linked 1% gelatin and FITC-labeled fibronectin to detect ECM degradation, and cells were incubated overnight in invadopodia medium for approximately 18 hours.

### 2.5 Immunofluorescence

For YAP and ROCK2 quantitation based on average pixel intensity, cell and nuclear areas were identified by immunostaining with an anti-p120-catenin rabbit primary polyclonal antibody (generated and gifted by Al Reynolds, Vanderbilt University) or rhodamine phalloidin (ThermoFisher) and Hoechst (ThermoFisher), respectively, in PAA invadopodia assays but with unlabeled fibronectin. YAP was identified using the same antibody for Western blotting as noted above, and ROCK2 was identified with a different anti-ROCK2 mouse monoclonal antibody (MilliporeSigma, Cat# WH0009475M1, RRID:AB_1843399). The relative nuclear ratio of YAP is a standard method that correlates with gene expression.^31,32^ As previously described, invadopodia were identified by immunostaining for F-actin and cortactin using rhodamine phalloidin and the 4F11 antibody (MilliporeSigma, Cat# 05-180, RRID:AB_309647), respectively.^20,21,29^ Invadopodia were counted manually, and ECM degradation was quantitated by thresholding for the loss of FITC signal under each cell.

### 2.6 Statistics

As previously described, statistics were performed using SPSS Software (IBM).^20,21,24,33-35^ All data were evaluated for normality with the Shapiro-Wilk or Kolmogorov-Smirnov test. Normal and non-parametric data were then analyzed by the Student’s t-test or a Kruskal-Wallis test, respectively, with significance at a p-value < 0.05.

## 3. RESULTS

While actomyosin contractility can facilitate the nuclear translocation of YAP, transcriptional activity can vary in different cell types due to phenotypic differences in mechanical responses to ECM rigidity.^6,23,31,36-44^ We have previously shown that ECM rigidity does promote actomyosin contractility by increasing pMLC levels in SCC-61 cells.^20,21^ Therefore, we first tested whether ECM rigidity also promotes nuclear accumulation of YAP given these established changes in cellular force generation. We performed invadopodia assays on soft and hard PAAs and measured the relative levels of YAP in the nucleus versus the cytoplasm of SCC-61 cells using quantitative immunofluorescence (Figures 1A-B). The ratio of these average intensity levels, which are independent of cell size, increased with the elastic moduli of the PAAs (i.e., from soft to hard) (Figure 1C). These results confirm an increase in nuclear YAP accumulation in response to ECM rigidity in SCC-61 cells.

**Figure 1.**
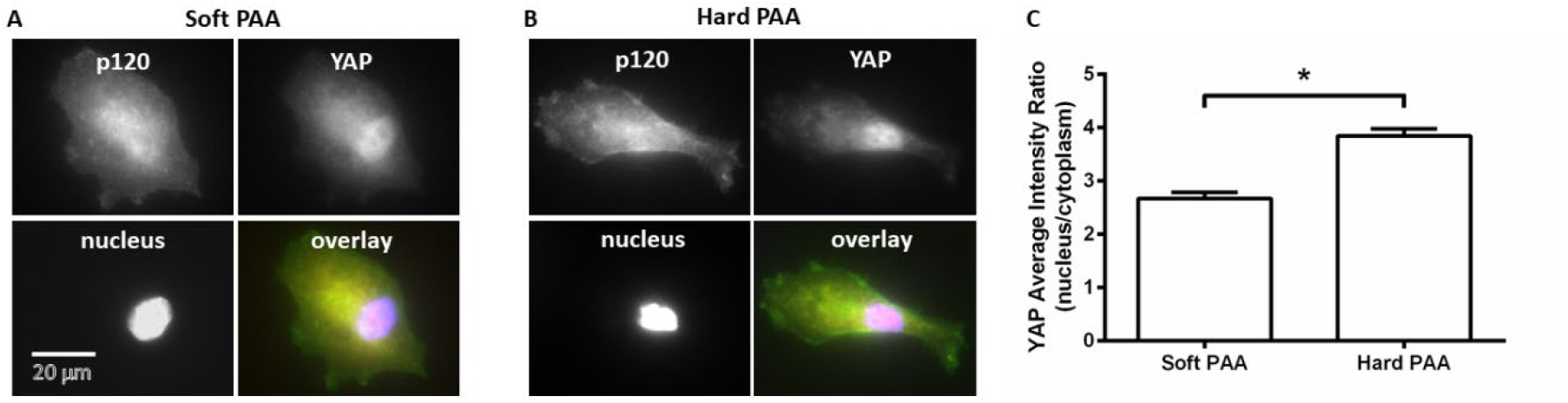
YAP nuclear translocation is regulated by ECM rigidity. Representative wide-field immunofluorescence images of the nuclear accumulation of YAP relative to the cytoplasm in SCC-61 cells on (A) soft and (B) hard PAA invadopodia assays. (C) Quantitative immunofluorescence based on average pixel intensities revealed an increase in nuclear YAP levels with increasing ECM rigidity. Data are shown as averages with standard error and * indicates p<0.05 for n=81-86 cells from 3 independent experiments.

YAP has recently been found to be a rigidity-dependent transcriptional regulator of ROCK2 expression in MCF-10 human breast cancer cells which also generate rigidity-dependent traction forces.^23,45^ Therefore, we also determined whether ROCK2 expression increased in SCC-61 cells with ECM rigidity using soft and hard PAAs and quantitative immunofluorescence (Figures 2A-B). Since ROCK2 is localized throughout the cell, we evaluated the average intensity across the cell body, and found an increase in ROCK2 expression in response to the elastic moduli of the PAAs when going from soft to hard substrates (Figure 2C). These results indicate that ROCK2 expression is also rigidity-dependent and suggest regulation by YAP activity.

**Figure 2.**
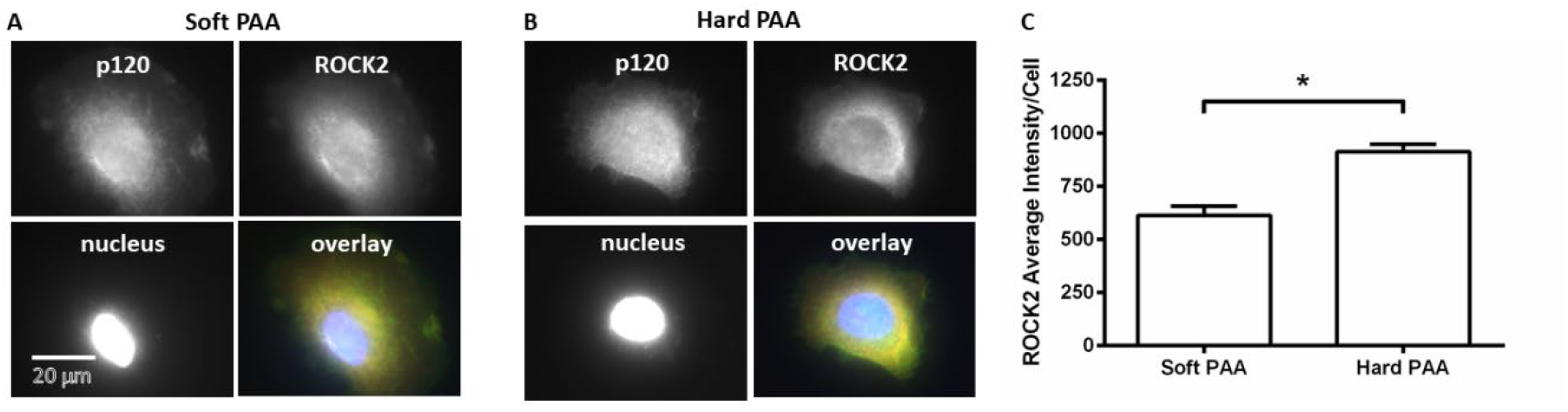
ROCK2 cellular levels are regulated by ECM rigidity. Representative wide-field immunofluorescence images of ROCK2 in SCC-61 cells on (A) soft and (B) hard PAA invadopodia assays. (C) Quantitative immunofluorescence based on average pixel intensities revealed an increase in ROCK2 cellular levels with increasing ECM rigidity. Data are shown as averages with standard error and * indicates p<0.05 for n=79-84 cells from 3 independent experiments.

To verify whether YAP regulates the levels of ROCK2 in SCC-61 cells, we examined both ROCK2 gene and protein expression using nested PCR and Western blotting, respectively, in response to the YAP inhibitor verteporfin, which inhibits YAP/TEAD interactions necessary for binding promoter sequences of target genes to initiate transcription (Figure 3).^26,27^ Verteporfin inhibited both the gene (Figure 3A) and protein (Figure 3B) expression of ROCK2 in SCC-61 cells. These results confirm that ROCK2 expression is dependent on YAP activity as a transcriptional coactivator.

**Figure 3.**
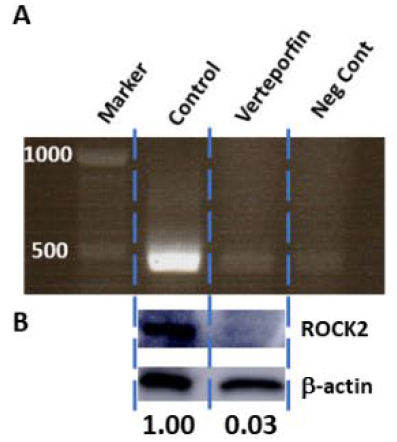
YAP regulates the expression of ROCK2. (A) Nested PCR and (B) Western blot revealed a decrease in ROCK2 gene expression and protein levels, respectively, after treatment with 20 μM verteporfin, an inhibitor of YAP/TEAD interactions which are necessary for the transcriptional activity of YAP.

Since we have now established a role for YAP in regulating ROCK2 which we have previously found to be important for invadopodia maturation,^21^ we then tested whether YAP also regulated this process and subsequent ECM degradation (Figure 4). To minimize the risk of any off-target effects caused by verteporfin, we specifically knocked down YAP using a single ON-TARGETplus siRNA (Figure 4A) which also inhibited the expression of ROCK2 (Figure 4B) in SCC-61 cells as confirmed by Western blots. We then performed invadopodia assays on the hard PAAs (Figure 4C) and found a significant decrease in ECM degradation (Figure 4D) and active (i.e., mature) invadopodia (Figure 4E). However, we found no difference in the number of total invadopodia (Figure 4F) which consist of both actively degrading and non-degrading or immature structures as denoted by yellow and white boxes in Figure 4C, respectively. Therefore, these results suggest that YAP regulates invadopodia maturation but not formation consistent with the activity of its downstream target ROCK2.

**Figure 4.**
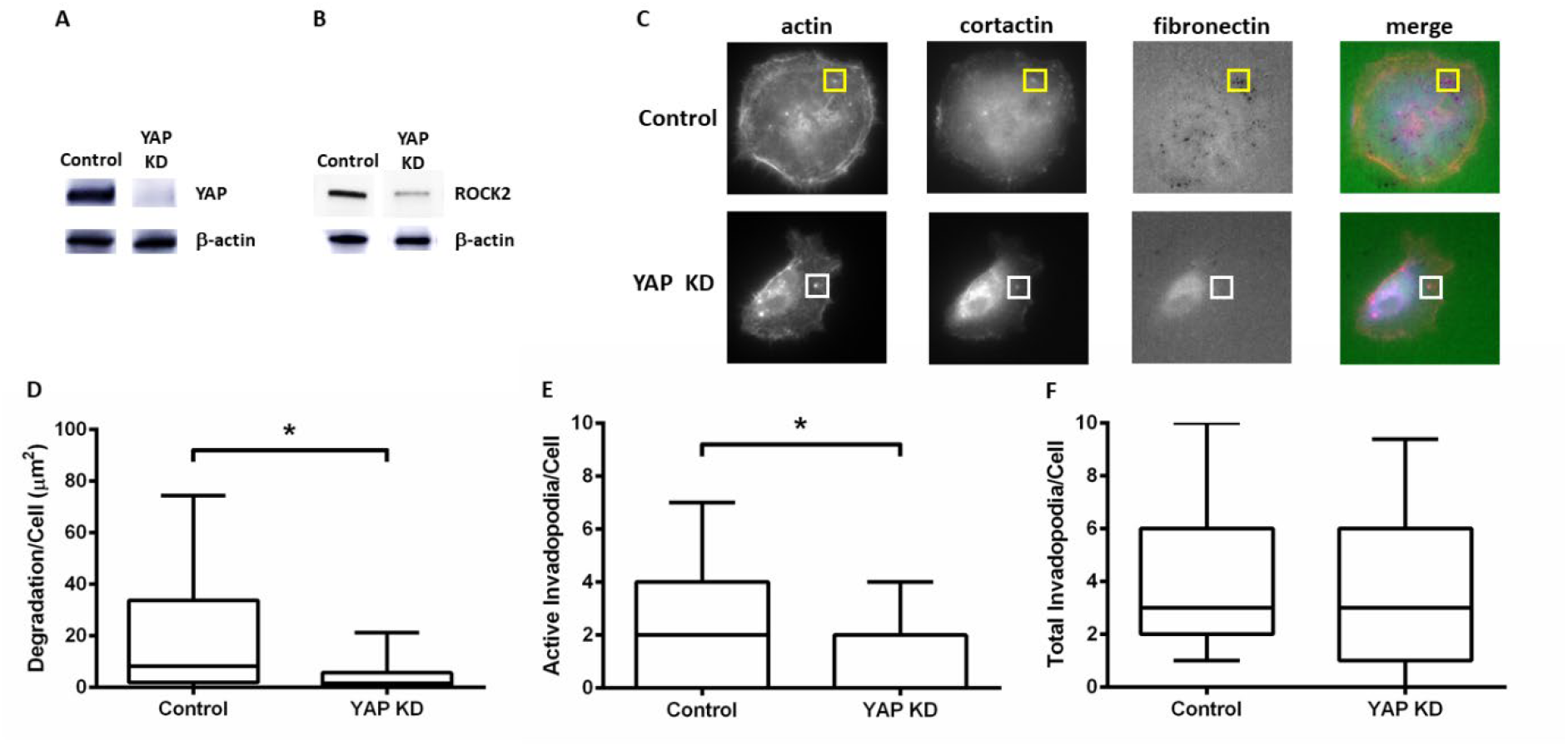
YAP regulates the formation of mature invadopodia and ECM degradation. (A) siRNA-mediated knockdown (KD) of YAP (B) inhibited the expression of ROCK2 in SCC-61 cells as indicated by Western blots. (C) Representative wide-field immunofluorescence images of non-targeting control and YAP KD in SCC-61 cells in hard PAA invadopodia assays. Immature or non-degrading invadopodia (white boxes) were identified by the colocalization of actin (red) and cortactin (blue) while mature or actively degrading invadopodia (yellow boxes) were further colocalized with black degraded areas lacking FITC-labeled fibronectin. (D) ECM degradation and (E) the number of mature or actively degrading invadopodia decreased with YAP KD while (F) total number of invadopodia (mature and immature) remained the same. Data are shown in (D-F) as box and whisker plots with black lines indicating medians and whiskers representing the 10th and 90th data percentiles and * indicates p<0.05 for n=45-51 cells from 3 independent experiments.

## 4. DISCUSSION

Mechanosensitive molecules such as ROCK that are activated in signaling pathways promoted by tumor biomechanical properties are emerging as significant factors in HNSCC and correlate with poor prognoses and reduced survival rates.^30,46-66^ YAP and its paralog WW domain containing transcription regulator 1 (TAZ) have been identified as critical intracellular mechanotransducers that can drive proliferation, growth, and metastasis in HNSCC.^8-13^ YAP and TAZ are often viewed as redundant, but functional differences are emerging due to an additional binding motif in YAP.^44,67-71^ Furthermore, only YAP has been correlated with poor clinical outcomes in HNSCC.^10-13^ Therefore, we chose to focus on YAP to determine whether this transcriptional coactivator played a role in invadopodia activity which correlates with invasive behavior and tumor progression.^14-16^ Overall, our results from this study suggest that YAP can mediate invadopodia maturation and ECM degradation through its regulation of ROCK2.

YAP localization and activity are mediated by several mechanisms, most notably through negative regulation by the Hippo pathway.^38,39^ However, this process can be overridden by ECM rigidity and force-dependent mechanisms.^70-72^ ECM rigidity can drive actomyosin contractility to promote the entry of YAP into the nucleus by stretching nuclear pores and thus activating the TEAD family of transcription factors for gene expression.^38^ Similar to other studies, we also found that an increase in ECM rigidity enhances the nuclear accumulation of YAP in SCC-61 cells which coincides with previously established trends in force generation using the same substrates.^20,21^ These increases in actomyosin contractility in response to ECM rigidity are consistent with other types of cancer cells that also make invadopodia including our previous work with breast cancer cells.^73-77^ Furthermore, the relative changes in nuclear translocation that we found in SCC-61 cells are similar to those in other studies that were sufficient to alter transcription, downstream signaling pathways, and resulting cellular phenotypes.^31,36-44,78^

Although YAP has been implicated in processes that enhance cell migration which is required for invasive behavior,^13,39,79^ a role for YAP in proteolytic activity which is necessary for breaching dense ECMs during invasion is unknown. We have previously shown that actomyosin contractility promotes invadopodia maturation and activity downstream of force generation through a ROCK2-dependent mechanism that affects actin filament stability via rigidity-dependent activation of LIMK.^20,21,77^ ROCK2 has recently been identified as a transcriptional target of YAP in MCF-7 breast cancer cells;^23^ however, YAP depletion in CAL51 breast cancer cells had no effect on ROCK2 gene expression.^39^ Since we found that YAP nuclear accumulation increased with ECM rigidity and is correlated with target gene expression, we first determined whether ROCK2 protein expression also increased with ECM rigidity. Not only did we find an increase in the cellular levels of ROCK2 in SCC-61 cells with ECM rigidity, but we also confirmed that ROCK2 protein and gene expression were inhibited by the YAP inhibitor verteporfin validating that YAP is a transcriptional regulator of ROCK2 in SCC-61 cells. These results are consistent with previous studies that have shown increased expression and/or nuclear accumulation of YAP in human HNSCC tumors which are more rigid than normal tissues.^8,11-13,56,80-82^

While YAP has been associated with HNSCC, we are unaware of any studies in which this transcriptional coactivator has been directly linked to invadopodia regulation despite the presence of the YAP gene in a common HNSCC amplicon.^10,83^ Given our findings that YAP regulates ROCK2 which we have previously shown to promote invadopodia maturation,^21^ we then determined whether YAP also impacted this process in SCC-61 cells. KD of YAP led to a decrease in ECM degradation and the number of mature or actively degrading invadopodia. However, we also found that YAP KD had no effect on the total number of invadopodia (mature and immature or nascent). These findings phenocopy the effects of ROCK2 inhibition on active invadopodia and proteolysis as previously reported.^21^ Therefore, our results indicate that YAP also promotes the formation of mature invadopodia downstream from actomyosin contractility.

## 5. CONCLUSION

In this study, we have shown that rigidity-dependent YAP localization promotes ROCK2 expression and regulates invadopodia maturation and activity. While further studies are required, our data suggest a means by which actomyosin contractility can promote YAP-mediated proteolytic invasion by affecting local actin dynamics at invadopodia in HNSCC cells. Actomyosin contractility is regulated by several other kinases besides ROCK including myosin light chain kinase (MLCK) and p21-activated kinase (PAK).^84-86^ However, PAK can have divergent effects on invadopodia, and we have recently reported that MLCK does not alter actomyosin contractility nor decrease invadopodia activity in HNSCC cells with siRNA inhibition.^21,25,87-91^ Our previous work has established a role for ROCK1, but not ROCK2, in regulating contractile forces via pMLC while only ROCK2 activated LIMK which is similar to the findings of another study regarding the ROCK isoforms in MDA-MD-231 breast cancer cells which we have also found to exhibit rigidity-dependent actomyosin contractility.^21,92^ In addition, YAP does not regulate ROCK1 transcription thereby preserving actomyosin contractility.^23^ Therefore, YAP may serve as a link between the activity of the ROCK isoforms to drive migration and proteolytic invasion of cancer cells. Furthermore, ROCK2 has also been found to enhance YAP nuclear translocation and transcriptional activity suggesting a positive feedback loop for potentiating invadopodia dynamics.^22,23,93,94^ Overall, these findings suggest a novel mechanism by which force-mediated transcriptional regulation may direct proteolytic activity by controlling the mechanically-induced formation of stable and actively degrading invadopodia.

## ACKNOWLEDGMENTS

This work was conducted in part using the resources of the Advanced Computing Center for Research and Education at Vanderbilt University.

